# Chloroplastic ROS bursts initiate salicylic acid biosynthesis in plant immunity

**DOI:** 10.1101/2024.08.26.609370

**Authors:** Charles Roussin-Léveillée, Méliane St-Amand, Philippe Desbiens-Fortin, Rosaëlle Perreault, Antoine Pelletier, Sabrina Gauthier, Faye Gaudreault-Lafleur, Isabelle Laforest-Lapointe, Peter Moffett

## Abstract

Chloroplasts are essential centers of signal integration and transduction in plants. They are involved in the biosynthesis of primary and specialized metabolites, including salicylic acid (SA), a key defense phytohormone synthesized via the conserved chorismate biosynthetic pathway. However, the identity of the signal(s) that ultimately triggers SA induction in chloroplasts upon perception of a biotic threat has remained elusive. Here, we provide evidence of a functional link between chloroplast-derived reactive oxygen species (cROS) and SA production. We observe that inhibiting ROS bursts generated from photosystem II during plant immune activation completely abrogates the induction of SA synthesis in response to immunity-inducing signals, without affecting SA-independent immune responses. Indeed, time course analyses show that the induction of SA marker genes parallels that of cROS production during an immune response. Consistent with this, preventing cROS induction is sufficient to nullify the immune protection normally conferred by activating immunity prior to an infection. Analyses of transcriptomes and photosynthetic efficiency show that two conserved effectors from the phytopathogen *Pseudomonas syringae*, HopM1 and AvrE1, redundantly disrupt photosynthesis and cROS bursts. These effects reduce SA accumulation and are mediated via the impact of HopM1 and AvrE1 in inducting host abscisic acid signaling. Our results suggest that a change in chloroplastic redox homeostasis induced by biotic stressors acts as an initiator of plant immunity through the production of SA, and that this response is targeted by conserved pathogen effector proteins.

## Introduction

Plant cells respond swiftly to external threats through the establishment of spatiotemporal, multi-layered, immune responses. Following the activation of pattern recognition receptors (PRRs) by immunogenic pathogen-associated molecular patterns (PAMP), an immune signaling cascade known as PAMP-triggered immunity (PTI) is induced^1^. PTI activation affects multiple physiological and biochemical processes, ultimately leading to pathogen resistance^1^. In the initial steps of PTI activation, PRR-co-PRR complexes associate with RESPIRATORY BURST OXIDASE HOMOLOGUE PROTEINS D and F (RBOHD/F) to generate reactive oxygen species (ROS) bursts in the apoplastic space, as well as calcium (Ca^2+^) channels to induce Ca^2+^ influx^1^. These early events at the plasma-membrane further lead to phosphorylation of mitogen-activated protein kinases (MAPK), biosynthesis of defense-associated phytohormones and a transcriptional reprogramming of the host^1^.

Defense-associated phytohormones, such as salicylic acid (SA), jasmonic acid and ethylene, serve as amplifiers of plant immune responses^2,3^. Multiple plant hormones associated with defense, including SA, are synthesized in the chloroplast^4,5^. As such, a multifaceted relationship exists between the plant photosynthetic center and the establishment of robust defense responses. Indeed, chloroplasts have been identified as being directly involved in reinforcing immune responses against phytopathogens ranging from bacteria, fungi, oomycetes and viruses^6–10^. Photoperiodic stress, such as excess light, causes damage to chloroplasts and triggers plant transcriptional responses reminiscent of a PTI response^11–13^. Furthermore, light is essential for activating immune signaling cascades to prevent pathogen infection^12,14^. Beyond their critical involvement in PTI, chloroplasts have also been shown to be essential actors in effector-triggered immunity (ETI)^4–7,9^, ETI is generally a stronger response compared to PTI and is induced by the recognition of proteins, known as effectors, that are delivered to the plant cytoplasm to promote disease.

Chloroplasts generate ROS bursts during immune response and it has been reported that chloroplasts are targeted by microbial effectors^15^. The function of these chloroplastic ROS (cROS) in plant immunity is still unclear, although their disruption has been shown to increase the virulence of a hypovirulent bacteria^15^. It has been suggested that cROS disruption by *Pseudomonas syringae pv. tomato* (*Pst*) DC3000 is mediated by the actions of *Pst* effectors on the host abscisic acid (ABA) biosynthesis program and that this contributes significantly to bacterial colonization^15^. We and others, have identified two conserved effectors from *Pst*, namely HopM1 and AvrE1, as being involved in the manipulation of the plant ABA biosynthesis and signaling and biosynthesis programs^16,17^. Here, we investigate the impact of HopM1 and AvrE1 on the disruption of photosynthesis and cROS bursts, as well as the implications of this disruption on microbial virulence.

Although cROS bursts have been observed in many studies in recent decades^18^, and are sometimes used as reporters of plant immune activation, their function in immunity has remained elusive. In this study, we provide evidence that cROS bursts are essential for SA-dependent immune functions and that their induction correlates with that of SA biosynthesis and signaling. Genetic and functional analysis show that cROS production occurs upstream of SA induction, whereas SA does not induce cROS bursts. Transcriptome and high-resolution phenotyping analyses showed that plants infected by WT *Pst*, but not by a *Pst* strain lacking the effectors HopM1 and AvrE1, have reduced photosynthetic functions, as well as a down-regulation of photosynthetic and chloroplast-encoded transcripts. Disruption of cROS by HopM1 and AvrE1 was dependent on their manipulation of the host ABA pathway, consistent with previously reported effects of these two effectors.

## Results

### Photosynthetic functions are required for immune function

We first investigated the importance of photosynthetic processes in the interaction between plants and *Pst*. To do so, Arabidopsis plants were treated with the photosystem II (PSII) inhibitor DCMU (3-(3,4-dichlorophenyl)-1,1-dimethylurea), also known as Diuron, in combination with the PAMP flg22, prior to a pathogen challenge. As expected, treatment with the immunogenic peptide flg22 resulted in an important protection against a subsequent infection by *Pst*, as indicated by infection symptoms and by bacterial counts. However, this protective effect was completely abrogated when co-infiltrated with DCMU (Fig. 1, A and B).

**Figure 1.**
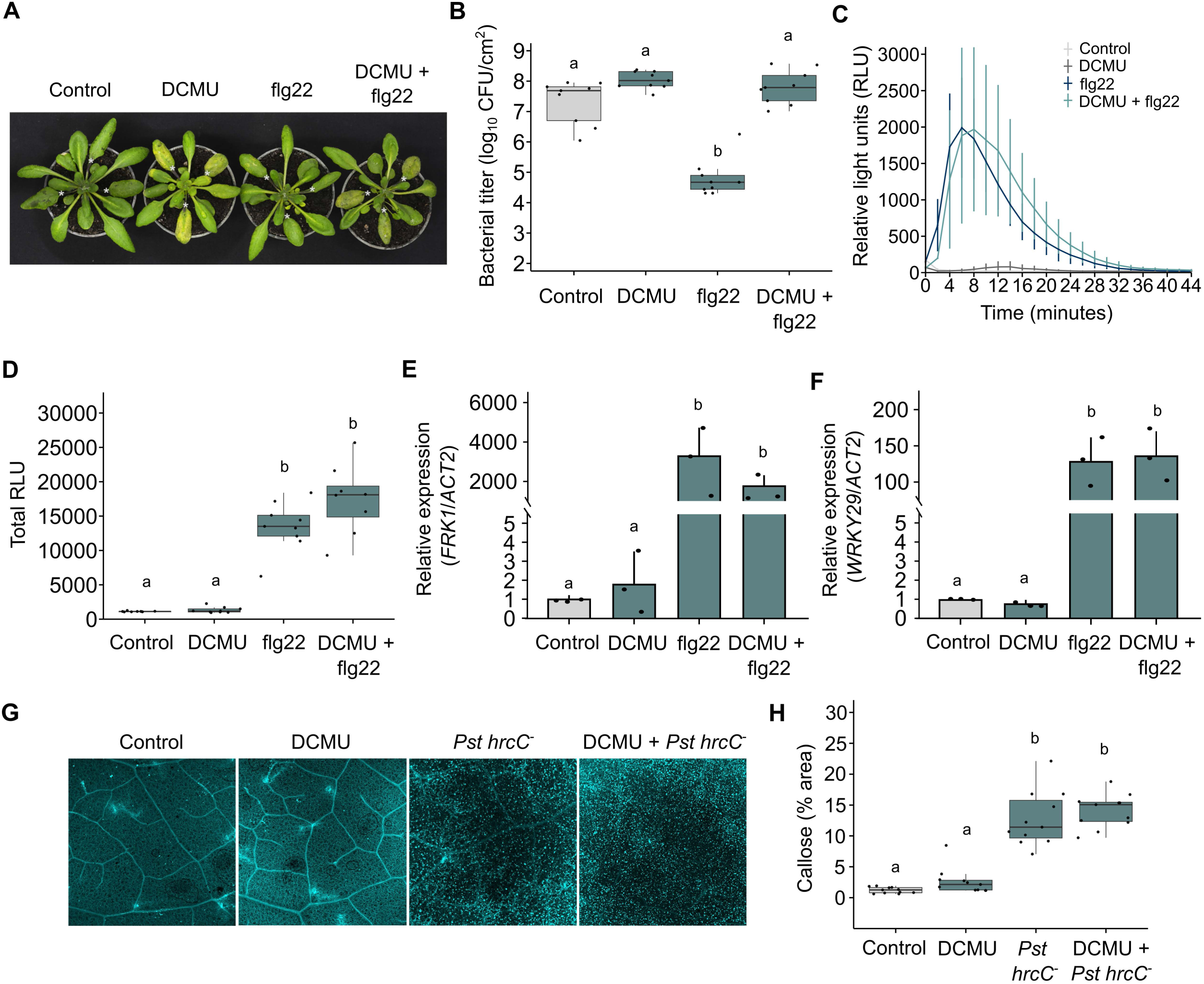
Inhibition of photosynthesis disrupts PAMP-mediated immune protection without preventing early immune signaling activation. (A) Leaves of four-week-old WT Arabidopsis plants were pre-treated by syringe-infiltration with 10 mM MgCl2 (control), DCMU (10 μM), flg22 (1 μM) or a combination of DCMU and flg22, as indicated. The same leaves were syringe-infiltration with *WT Pst* (5 x 10^5^ CFU/ml) 24 hours after the initial treatment. Plants were photographed three days post-infection (dpi). (B) Bacterial titers measured in leaves treated as in (A) at 3 dpi. (C) Arabidopsis plants were treated as in (A), followed by measurement of reactive oxygen species (ROS) measured as relative light units (RLU) over 44 min (n = 9 leaf discs/treatment). (D) Quantification of total RLU measured from the experiment in (C). (E-F) Arabidopsis plants were treated as in (A), followed by quantification of the relative expression of *FRK1* (E) or *WRKY29* (F) by qRT-PCR with *ACT2* acting as a reference gene. (G) Leaves of four-week-old Arabidopsis plants were syringe-infiltrated with 10 mM MgCl2 (control), DCMU (10 μM), *Pst hrcC*^-^ (1×10^8^ CFU/ml) or a combination of the bacterial inoculum with DCMU, as indicated. Leaf punches were harvested at 24 hpi and subjected to callose staining with aniline blue, followed by confocal microscopy. (H) Callose deposits were quantified as a percentage of total image area (% area) as determined by ImageJ from the samples described in (G). All experiments were repeated at least three times with similar outcomes. All figures containing datapoints were subjected to a Kruskal-Wallis test to compare the different treatments. Different letters indicate statistical significance.

Plasma-membrane activation of plant immune responses rapidly leads to apoplastic ROS bursts by the NADPH oxidases RBOHD/F^19^. We evaluated whether a treatment with DCMU would result in reduced apoplastic ROS potentiation triggered by flg22, which would suggest a dysfunction in early immune responses. We found that a co-treatment of flg22 with DCMU did not alter apoplastic ROS burst at the plasma membrane (Fig. 1, C and D). This suggests that DCMU does not significantly affect the PRR-coPRR complex from being activated and associating with RBOHD/F.

We next evaluated whether DCMU treatment impacts early signal transduction in the nucleus and prevents early immune-related genes from being expressed. We found no statistical differences in the expression levels of two early immune marker genes, *FRK1* and *WRKY29,* upon treatment with flg22 alone or in combination with DCMU (Fig. 1, E and F). Activation of early immune signaling commonly leads to cell-wall reinforcement through callose deposition^1^. We therefore evaluated whether a lack of callose deposition could explain the reduced immune protection by a flg22 pre-treatment, when co-treated with DCMU. We observed similar levels of callose deposition triggered by flg22, in the absence or in the presence of DCMU, at 24 hours post treatment (Fig. 1, G and H). Together, these results suggest that the abrogated immune response in the presence of DCMU does not seem to be caused by a defect in early immune signaling events or by a reduction in cell-wall reinforcement.

We also investigated whether DCMU could prevent ETI responses. High dose inoculum (1 x 10^8^ CFU/ml) of *Pst* strains carrying the avirulence genes AvrRpm1 or AvrRps4 induced ETI-associated hypersensitive responses in the presence or absence of DCMU (fig. S1A). This suggests that ETI-associated cell death is not affected by DCMU treatment under these conditions. However, a clear increase in growth of avirulent bacteria was observed when lower doses of bacterial inoculum (1 x 10^5^ CFU/ml) were co-infiltrated with DCMU compared to control conditions (fig. S1B). This suggests that inhibition of PSII results in suppression of ETI-mediated bacterial growth inhibition.

### Chloroplastic ROS production is conserved in plant immunity and depends on photosynthesis

Production of cROS has been reported following the perception of certain bacteria and in plants subjected to abiotic stresses^4,11,20^. Since DCMU disrupts photosystem II (PSII) functions, and prevents cROS bursts following the perception of a type-III secretion system mutant of *Pst*^15^, we investigated the general involvement of cROS bursts in mediating plant immunity. We thus tested whether different PTI-inducers, including the PAMPs flg22 and elf18, and the damage-associated molecular pattern (DAMP) Pep1, trigger cROS production. This was assessed using the dye H_2_DCF-DA, which can be visualized by microscopy in the presence of ROS. We found that all three peptides triggered strong cROS production four hours after syringe-infiltration in Arabidopsis leaves (Fig. 2, A and B). This suggests cROS production is triggered by multiple inducers of PTI responses.

**Figure 2.**
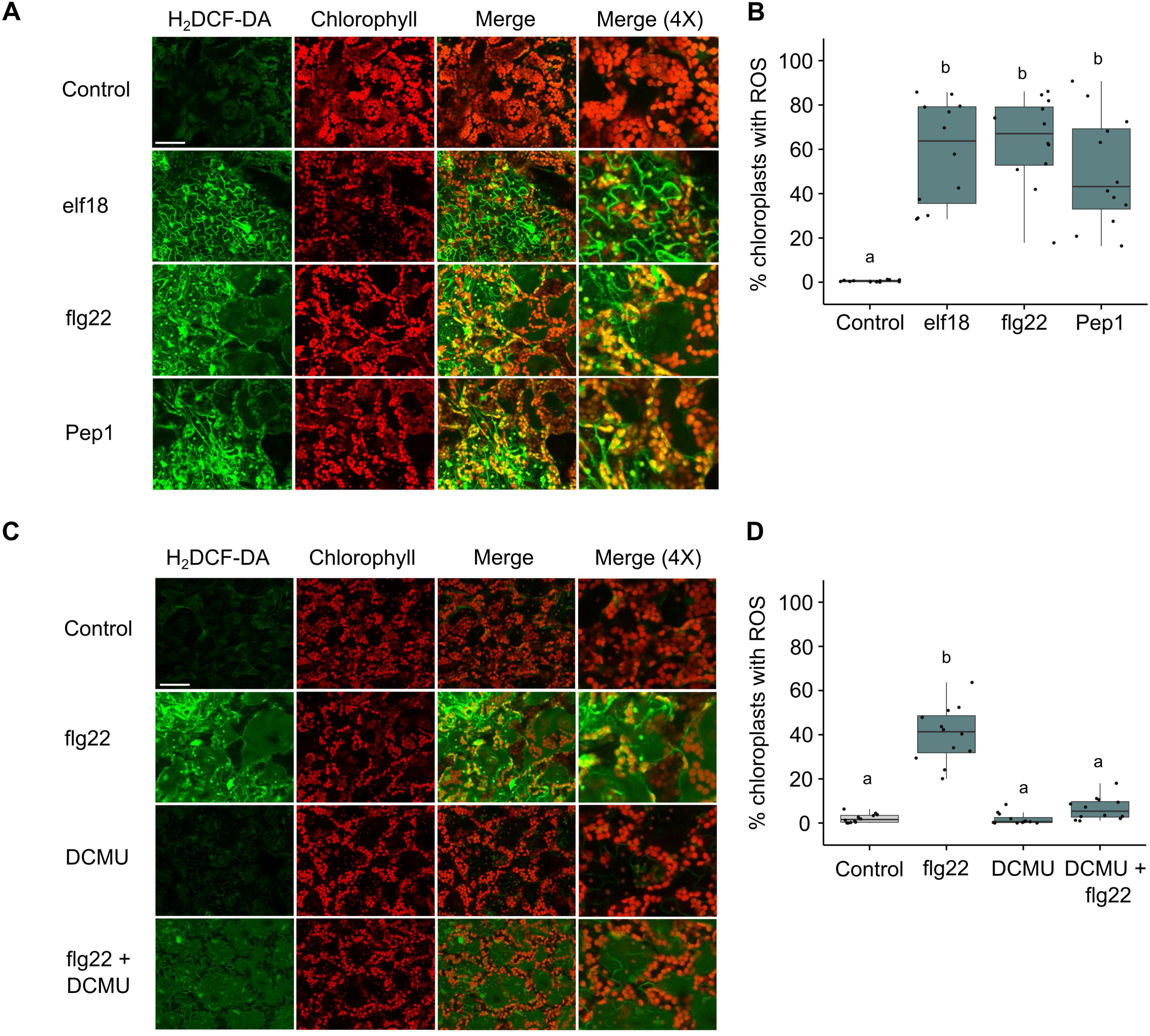
Immune stimulation induces chloroplastic ROS in a photosystem-dependent manner. (A) Leaves of four-week-old WT Arabidopsis plants were syringe-infiltrated with either 10 mM MgCl_2_ (control), flg22 (1 μM), elf18 (1 μM) or pep1 (1 μM), as indicated. Five hours later, leaves were infiltrated with H_2_DCF-DA and leaf punches immediately harvested and mounted on microscopy slides, followed by visualization by confocal microscopy. Panels display channels detecting ROS production (H_2_DCF-DA), the presence of chloroplasts (chlorophyll), as well as merged images of the two (merge). Bars represent 50 μM. (B) Quantification of cROS detected in (A), presenting the number of cROS-positive chloroplasts as a percentage of total chloroplasts in images (% chloroplasts with ROS). Image analysis was performed by using the cell counter function of ImageJ after background removal. (C) Plant leaves were treated as in (A), but with initial infiltrations of either MgCl_2_ (10 mM), flg22 (1 μM), DCMU (10 μM) or a combination of flg22 and DCMU, as indicated. (D) Quantification of cROS found in (C) as in (B). All experiments were repeated at least three times with similar outcomes. All figures containing data were subjected to a Kruskal-Wallis test to compare the different treatments. Different letters indicate statistical significance.

It has been reported that a hypovirulent *Pst* strain results in cROS production^15^, and our results indicate that PAMP or DAMP treatment induces a similar response. Thus, we have used flg22 as a representative treatment to induce PTI responses in the absence of any unforeseen effects of toxin production or other unknown confounding factors in promoting, or inhibiting, cROS bursts. Indeed, we found that flg22-triggered cROS bursts were largely suppressed upon a co-treatment with DCMU (Fig. 2, C and D), thus establishing an assay to investigate the function of cROS bursts.

### Timing of chloroplastic ROS production correlates with salicylic acid biosynthesis

Since DCMU abrogates flg22-mediated immune protection against pathogen infection without affecting early PTI signaling events, we sought to evaluate the impact of DCMU on the chain of events downstream of these early responses. We investigated the timing of cROS burst induction after immune elicitation with flg22. We found that cROS are detectable within 3-4 hours after flg22 treatment (Fig. 3, A and B). This could imply that DCMU prevents immune responses that start around this timepoint. The induction of salicylic acid synthesis and signaling occurs after the first wave of transcriptional responses to immunity elicitation^1^ and directly amplifies immune responses^2,3^. We found a strong correlation between the timing of cROS production and the induction of the SA biosynthesis marker gene *ISOCHORISMATE SYNTHASE 1* (*ICS1)*. This was quickly followed by the induction of the highly SA-sensitive reporter gene *PATHOGENESIS-RELATED 1* (*PR1)*, which implies SA signaling activation (Fig. 3, C and D). Thus, the timing of the events in our results are consistent with a link between cROS bursts and SA induction, which is initiated via the chloroplast-localized chorismate pathway.

**Figure 3.**
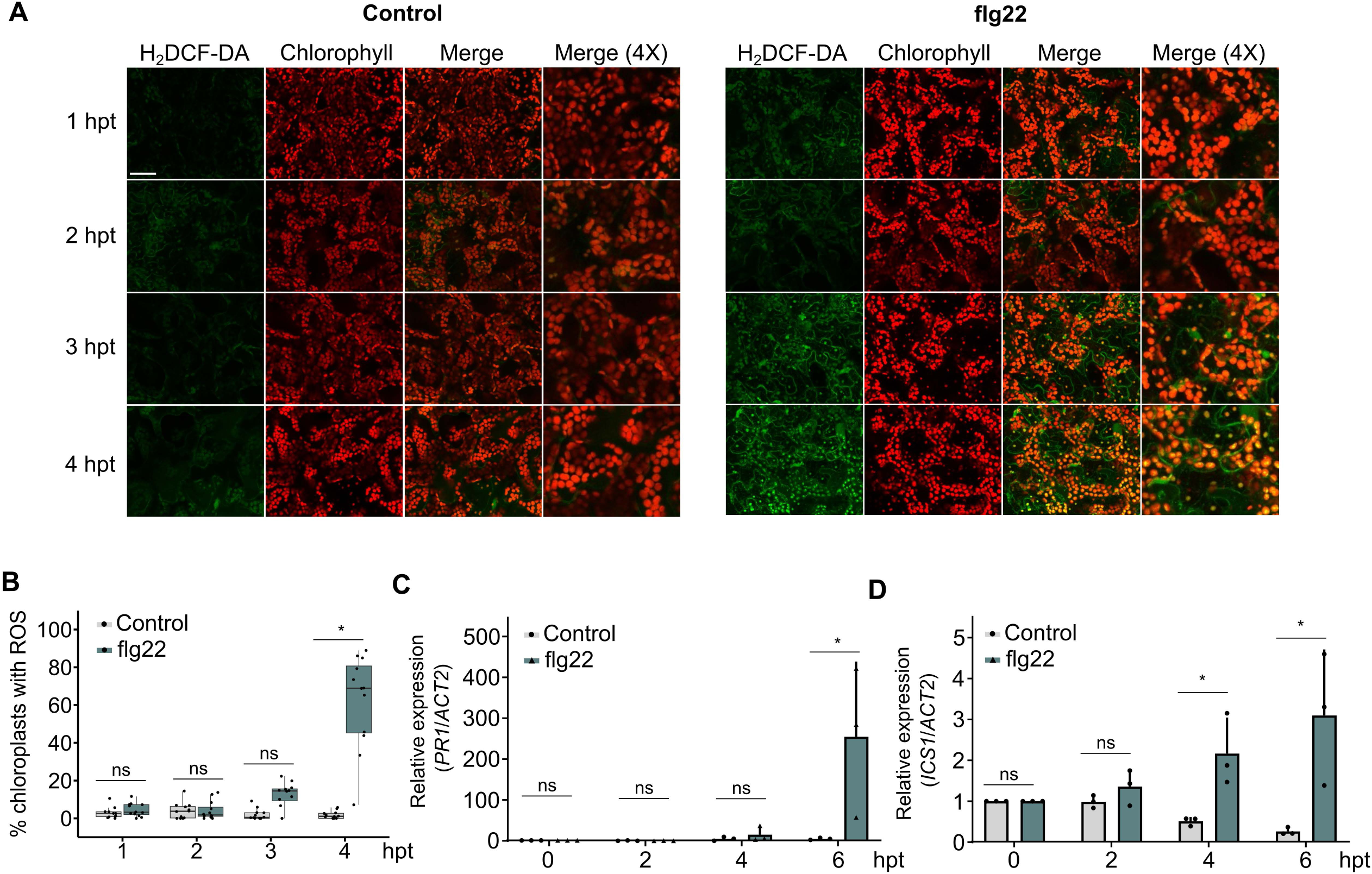
Timing of chloroplastic ROS production correlates with the onset of salicylic acid signaling. (A) Leaves of four-week-old WT Arabidopsis plants were syringe-infiltrated with either 10 mM MgCl_2_ (control) or flg22 (1 μM), as indicated. Leaves were subsequently infiltrated with H_2_DCF-DA at the indicated time points following initial treatment (hours post-treatment; hpt) and leaf punches immediately taken and mounted on microscopy slides. Panels display ROS production (H_2_DCF-DA), the presence of chloroplasts (chlorophyll) as well as a merge of the two (merge). Bars represent 50 μM. (B) Quantification of cROS detected in (A), presenting the number of cROS-positive chloroplasts as a percentage of total chloroplasts in images (% chloroplasts with ROS). Image analysis was performed by using the cell counter function of ImageJ after background removal. (C-D) Arabidopsis leaves were treated as in (A), followed by quantification of the relative expression of *PR1* (C) or *ICS1* (D) by qRT-PCR, at the indicated time points (hpt), with *ACT2* acting as a reference gene. Multiple Student’s t-test were performed in 3B-D to compare control to flg22 treatments. A * symbol indicates statistical significance (p < 0.05), while ns = not significant. All experiments were repeated at least three times with similar outcomes.

### Chloroplastic ROS induction is essential for PAMP-triggered SA biosynthesis

To test a potential link between cROS bursts and the initiation of SA biosynthesis, we evaluated the induction of expression of *ICS1* and *PR1* by flg22 treatment in plants co-treated or not with DCMU at 24 hours post-treatment. Whereas flg22 treatment results in a strong induction of expression of both *ICS1* and *PR1*, this induction is dramatically reduced when flg22 is co-infiltrated with DCMU (Fig. 4, A and B). Further supporting the impact of DCMU on SA pathways, we quantified SA, and its glycosylated storage form SAG, by UPLC-MS in Arabidopsis plants treated with either flg22 or DCMU, or a combination of the two. We found that DCMU repressed flg22-induced SA accumulation (Fig. 4, C and D). Inhibiting photosynthesis/cROS via application of DCMU did not prevent the induction of *PR1* expression when SA was applied exogenously in combination with DCMU (Fig. 4E), suggesting that DCMU only affects SA biosynthesis, and not SA signaling.

**Figure 4.**
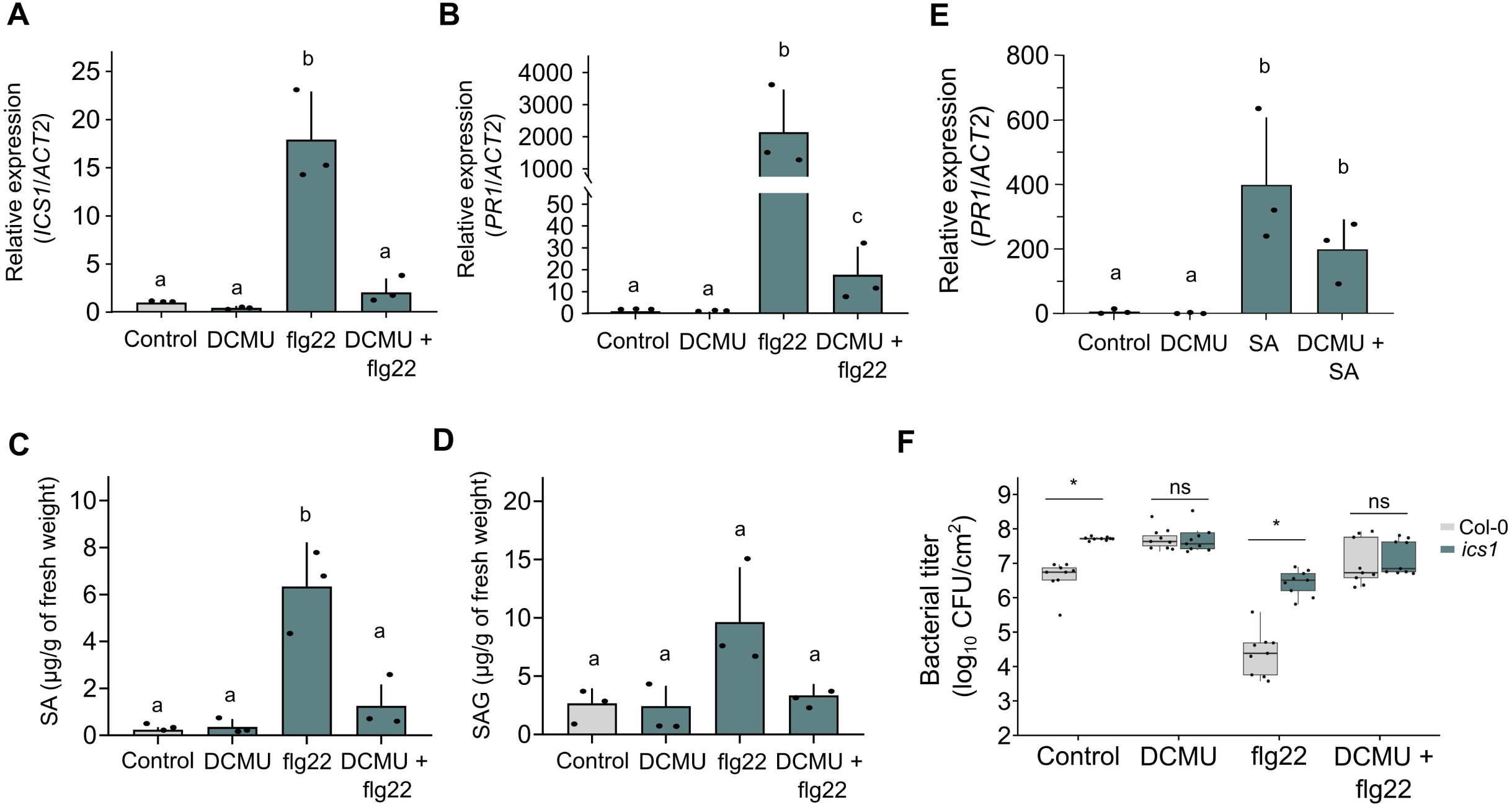
Inhibition of chloroplastic ROS induction/photosynthetic activities blocks PAMP-triggered SA accumulation and signaling. (A-B) Leaves of four-week-old WT Arabidopsis plants were syringe-infiltrated with 10 mM MgCl_2_ (control), flg22 (1 μM), DCMU (10 μM) or a combination of flg22 and DCMU, as indicated. At 24 hours post treatment (hpt), the relative expression levels of *ICS1* (A) and *PR1* (B) were determined by qRT-PCR with *ACT2* acting as a reference gene. (C-D) Arabidopsis leaves were treated as in (A-B) and harvested at 24 hpt, followed by quantification of salicylic acid (SA) (C) or salicylic acid beta-glucoside (SAG) (D). (E) Leaves of four-week-old WT Arabidopsis plants were syringe-infiltrated with 10 mM MgCl_2_ (control), DCMU (10 μM), SA (0.5 μM) or a combination of DCMU and SA, as indicated. At 24 hours post treatment (hpt), relative expression of *PR1* was determined by qRT-PCR with *ACT2* acting as a reference gene. (F) WT Arabidopsis and the SA biosynthesis mutant *ics1* were pre-treated with 10 mM MgCl_2_ (control), flg22 (1 μM), DCMU (10 μM) or a combination of flg22 and DCMU. Plants were syringe-infiltrated with *Pst* DC3000 (1 × 10^5^ CFU/ml) 24 hours later. Bacterial titers were measured at 3 days post infiltration with bacteria (*n* = 3 biologically independent replicates). Statistical analyses were performed as followed. Kruskal-Wallis tests were performed for 4A, 4B and 4E. ANOVA followed by a Tukey HSD test was performed for 4C and 4D. Different letters indicate statistical significance. Multiple Student’s t-test were performed in 5F to compare control to DCMU treatments. A * symbol indicates statistical significance (p < 0.05), while ns = not significant. All experiments were repeated at least three times with similar outcomes.

To further evaluate whether SA biosynthesis occurs downstream of cROS production, we investigated flg22-mediated cROS bursts in Arabidopsis *ics1* mutant plants, which cannot produce SA. We found that cROS bursts were induced upon treatment with flg22 in the *ics1* mutant, similar to WT plants (Fig. S2A and S2B). At the same time, SA treatment alone did not induce cROS (Fig. S2C and S2D), suggesting that SA biosynthesis is downstream of cROS production. We further evaluated the immune protection provided by flg22 in Arabidopsis *ics1* mutant plants. The immune protection conferred by a pre-treatment with flg22 was entirely lost in the *ics1* mutant, with levels of bacterial growth similar to those observed in untreated WT plants (Fig. 4F). WT plants treated with DCMU showed bacterial growth similar to *ics1* mutants, suggesting that blocking cROS production has an effect akin to abrogating SA production. Pre-treatment of WT, but not *ics1* mutants, with flg22 resulted in a protective effect against *Pst*. However, no significant differences in bacterial growth were observed between Col-0 and *ics1* mutant plants treated with a combination of flg22 and DCMU, further indicating that blocking cROS production has an effect comparable to abrogating SA production (Fig. 4F). These observations further support the role for cROS as initiators of SA biogenesis, as only this aspect of immunity appeared to be affected by DCMU. At the same time, if has been reported that *Pst* induces ABA production in Arabidopsis chloroplasts while disrupting photosynthesis, cROS bursts and SA production^15–17^, suggesting that not all phytohormone biosynthesis are affected by a reduction in photosynthesis.

### The conserved bacterial effectors HopM1 and AvrE1 disrupt photosynthesis for virulence

Considering the importance of the photosynthesis, cROS and SA in plant immunity, it is likely that this module is targeted by plant pathogens. Indeed, *Pst* has previously been reported to disrupt photosynthesis via type III secreted effector proteins, and this activity appears to depend on pathogen manipulation of the host ABA pathway^15^. We have previously reported the effects of water-soaking effectors, HopM1 and AvrE1, and found that these effectors induce ABA synthesis and signaling, resulting in the creation of an aqueous apoplast^16^. Using several approaches, we analyzed photosynthesis-related parameters in Arabidopsis inoculated with WT *Pst*, as well as single and double mutants of *HopM1* and *AvrE1*. To investigate photosynthetic output, we first measured the maximum dark-adapted quantum efficiency of PSII (*F_v_*/*F_m_*), which is commonly used to evaluate how a given stress affects PSII in a dark-adapted state^21^. Leaves inoculated with WT *Pst*, as well as the *Pst* single mutants for HopM1 or AvrE1, displayed reduced *F_v_*/*F_m_* at 24 hpi compared to non-infected plants (Fig. 5, A and B). However, the *Pst hopM1^-^/avre1^-^* (*h^-^/a^-^*) double mutant showed no reduction in *F_v_*/*F_m_* compared to control plants (Fig. 5B). We also evaluated the nonphotochemical chlorophyll fluorescence quenching (NPQ) in the same inoculated leaves, a process whereby excess absorbed light energy is dissipated as heat^21^. In agreement with a previous report^15^, we found that WT *Pst* leads to a suppression of NPQ. Interestingly, this process seems to be independent of the secretion of HopM1 and AvrE1, as the NPQ was similar in leaves inoculated with WT *Pst* and *Pst h^-^/a^-^*(Fig. 5C). These observations suggest that, while HopM1 and AvrE1 suppress photosynthesis, as determined by *F_v_*/*F_m_* quantification, other effectors are also involved in manipulating chloroplast functions, as determined by NPQ quantification.

**Figure 5.**
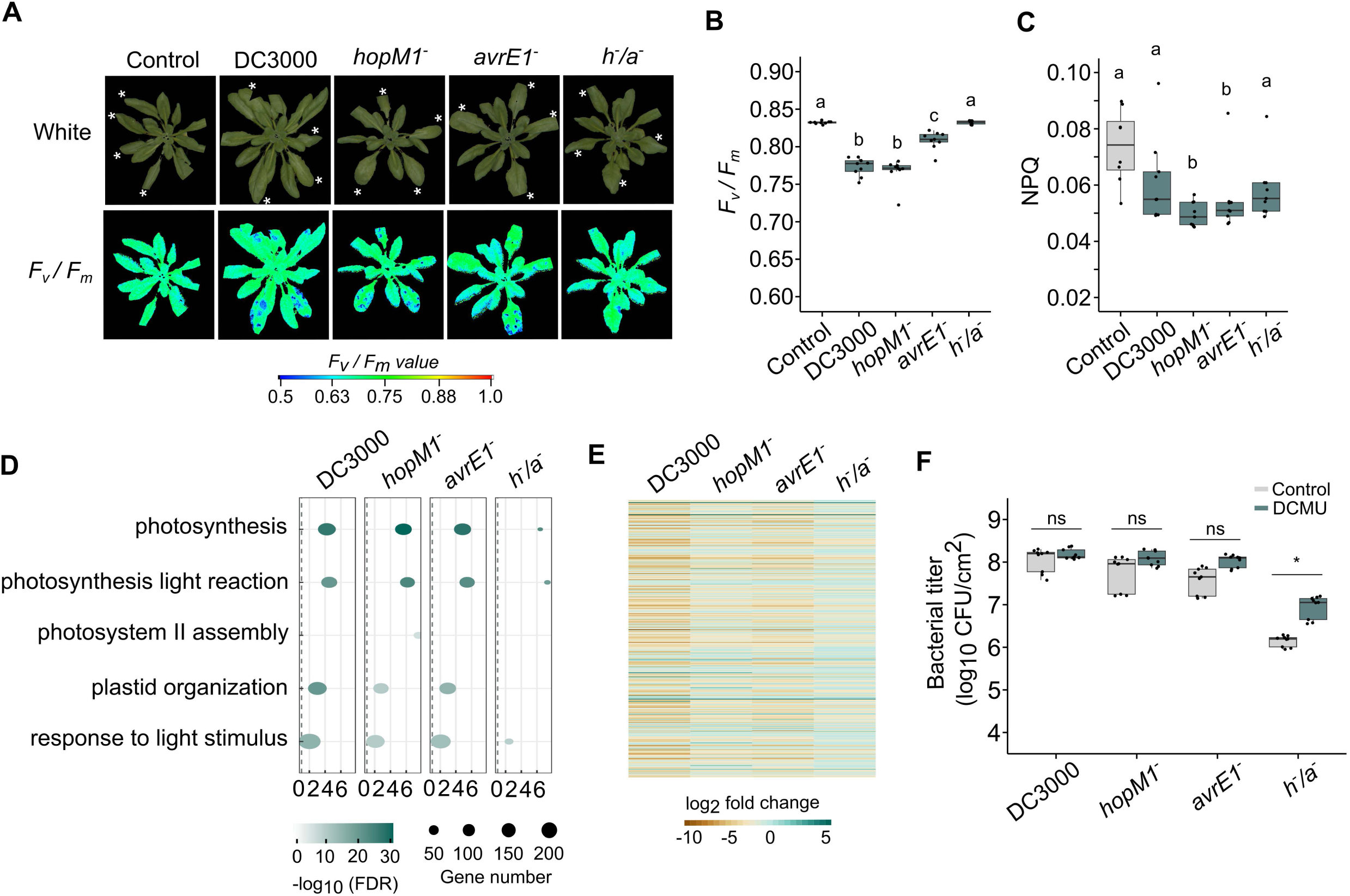
Pst water-soaking effectors reduce host photosynthetic functions. (A) Four-week-old WT Arabidopsis plants were infiltrated with either 10 mM MgCl_2_ (control) or bacterial strains (1 × 10^8^ CFU/ml), including *Pst* DC300, or mutant strains lacking either AvrE1, HopM1 or both (*h^-^/a^-^*). Plants were imaged at 24 hpi. Upper panel images represent RGB photos of the plants illuminated with white light (white). An image filter was applied to isolate plant tissues from background pixels. Lower panel images represent false-color analysis of the *F_v_/F_m_* values. (B) Quantification of *F_v_/F_m_* values obtained in (A) by averaging pixels. (C) Quantification of NPQ values obtained from a NPQ protocol on the same system as in (A) by averaging pixels from infiltrated leaf areas. (D) Gene ontologies associated with photosynthetic processes identified in previously published RNA-seq datasets (Roussin-Léveillée et al., 2022) from Arabidopsis leaves infiltrated with the indicated *Pst* strains (1 × 10^6^ CFU/ml) at 36 hpi. (E) Heatmap representing expression differences of genes associated with the gene ontologies in (D). (F) WT Arabidopsis and *aba2-1* mutant plants were syringe-infiltrated with *Pst* DC3000, *Pst hopM1^-^*, *Pst avrE1^-^* or *Pst h^-^/a^-^* (1 × 10^5^ CFU/ml), in the absence or presence of DCMU (10 μM) at 3 dpi, as indicated, followed by quantification of bacterial titers at 3 dpi. Statistical analyses were performed as followed. Kruskal-Wallis tests were performed for 5B and 5C. Different letters indicate statistical significance. Multiple Student’s t-test were performed in 5F to compare control to DCMU treatments. A * symbol indicates statistical significance (p < 0.05), while ns = not significant. All experiments were repeated at least three times with similar outcomes.

Next, we questioned whether HopM1 and AvrE1 affect the expression of photosynthesis-related genes. We analyzed the expression of photosynthesis-related gene ontologies (GO) of our previously reported transcriptomic data from Arabidopsis plants infected with WT *Pst*, as well as the single and double mutants for *HopM1* and *AvrE1*^16^. This analysis showed that HopM1 and AvrE1 cause a decrease in expression of a large number of nuclear-encoded genes associated with photosynthesis (Fig. 5D and E). Likewise, chloroplast-encoded genes are also negatively impacted by the action of these two water-soaking effectors (fig. S3). These results establish HopM1 and AvrE1 as important actors involved in the apparent suppression of photosynthesis during *Pst* infection in Arabidopsis.

To evaluate the importance of photosynthesis inhibition for HopM1 and AvrE1-mediated virulence, we investigated the impact of DCMU treatment on the virulence of *Pst* single and double mutant strains for both effectors. We found that, while a co-treatment of DCMU with WT *Pst*, *Pst hopM1^-^* or *Pst avrE1^-^* did not significantly affect bacterial growth compared to control, DCMU enhanced *Pst h^-^/a^-^* growth significantly (Fig. 5F). Interestingly, the growth of *Pst h^-^/a^-^* in combination with DCMU did not restore bacterial populations to the level observed with WT *Pst* in control conditions, suggesting that the impact of HopM1 and AvrE1 on *Pst* virulence is not solely dependent on their effects on photosynthesis. This lack of complete restoration of virulence could be explained by the inability of mutant *Pst* to induce water-soaking lesions, in either the absence or presence of DCMU (fig. S4A and S4B). Considering the level of conservation of AvrE1 across plant phytopathogens, it is likely that these observations could be mediated by AvrE homologs/orthologs in other pathosystems^22^.

### Water-soaking effectors disrupt cROS production to reduce salicylic-acid mediated defense responses

A connection seems to exist between host photosynthesis and the cROS production in plant immunity (Fig. 2C). Since HopM1 and AvrE1 cause a reduction in photosynthesis, we evaluated whether they also disrupt cROS production. We found that a *Pst h^-^/a^-^* mutant was impaired in its ability to suppress cROS production compared to *Pst* WT (Fig. 6, A and B). Interestingly, the secretion of either HopM1 or AvrE1 was largely sufficient to reduce cROS bursts during an infection, although not with the same level of consistency as that observed with *Pst* WT (Fig. 6, A and B). This is likely caused by the fact that single effector mutants are less efficient in manipulating host ABA responses^16,17^, which may result in greater variation in ABA concentrations and consequently lead to variation in cROS production. Importantly, the difference in cROS burst intensity between single and double *Pst* mutants for HopM1 and AvrE1 was not caused by a difference in bacterial population levels, as the high inoculum (1 x 10^8^ CFU/ml) used in these experiments allows for similar growth between the different *Pst* strains (Fig. 6C). As HopM1 and AvrE1 are not chloroplast-localized effectors, their impact on cROS is likely indirect and via their manipulation of the host ABA pathway.

**Figure 6.**
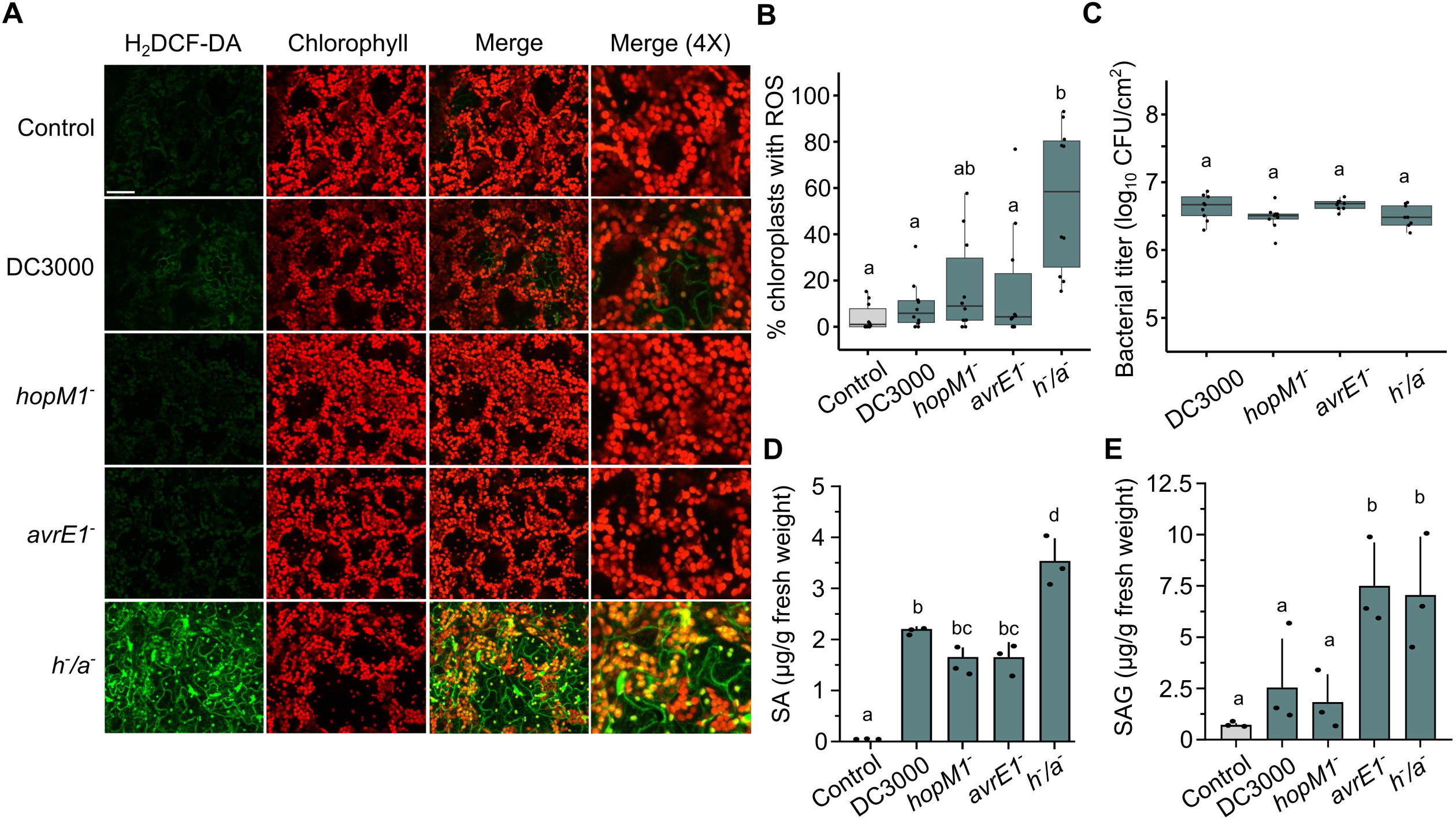
Water-soaking effectors inhibit chloroplastic ROS production to dampen the SA immune module. (A) Four-week-old WT Arabidopsis leaves were syringe-infiltrated with either 10 mM MgCl_2_ (control) or the indicated *Pst* strains (1 × 10^8^ CFU/ml). Twenty-four hours later, leaves were infiltrated with H_2_DCF-DA and leaf punches were immediately harvested and mounted on microscopy slides, followed by visualization by confocal microscopy. Panels display channels detecting ROS production (H_2_DCF-DA), the presence of chloroplasts (chlorophyll), as well as merged images of the two (merge). Bars represent 50 μM. (B) Quantification of cROS detected in (A), presenting the number cROS-positive chloroplasts as a percentage of total chloroplasts in images (% chloroplasts with ROS). Image analysis was performed by using the cell counter function of ImageJ after background removal. (C) Arabidopsis plants were treated as in (A), followed by quantification of bacterial titers at 24 hpi. (D-E) Arabidopsis plants were treated as in (A), followed by quantification of salicylic acid (SA) (D) or salicylic acid beta-glucoside (SAG) (E) by UPLC-MS at 24 hpi. All experiments were repeated at least three times with similar outcomes. Statistical analyses were performed as followed. For 6D, an ANOVA followed by a Tukey HSD test was performed. For 6E, a Kruskal-Wallis test was performed. Different letters indicate statistical significance (p < 0.05).

We also evaluated the effects of HopM1 and AvrE1 on SA accumulation in Arabidopsis plants. While SA and SAG still accumulated in plants challenged with WT *Pst* or single effector mutants, plants infected with *Pst h^-^/a^-^* accumulated at least twice as much SA and SAG (Fig. 6, D and E). We have previously demonstrated that a photoperiodic stress can prevent pathogen-induced water-soaking lesions by increasing SA levels^12^. We reasoned that, if DCMU indeed blocked SA accumulation during an infection, *Pst* would be able to induce water-soaking lesions under conditions of photoperiodic stress if co-infiltrated with DCMU. Consistent with our previous results^12^, we found that under a 24-hour constant light (CL) period, WT *Pst* did not induce water-soaking (fig. S5A). However, in plants inoculated with *Pst* and DCMU, water-soaking lesions were fully restored, resembling those observed in plants grown under a regular light (RL) photoperiod of 12 hours light/dark (fig. S5A). Bacterial growth in Arabidopsis WT plants under CL treated with DCMU were also similar to those found under RL, with or without a DMCU treatment (fig. S5B). This suggests that the resistance conferred by a CL treatment can be abrogated by DCMU, most likely via its disruption of SA production. Together these observations agree with a previous study suggesting that effectors located in the conserved effector locus of *Pst*, which includes HopM1 and AvrE1, inhibit SA-mediated basal immunity^23^. Our findings now suggest that HopM1 and AvrE1 promote bacterial pathogenesis by suppressing cROS production, thereby reducing SA-mediated functions in plant immunity.

### The missing link between SA and ABA pathways converges on cROS biogenesis

We and others have previously shown that *Pst* cannot create an ideal water-rich niche in the apoplast of Arabidopsis *aba2-1* mutant plants, which cannot produce ABA^16,17^. It has also been suggested that *aba2-1* mutants exhibit constitutively activated immune responses^24^. Here, we evaluated the intensity of this constitutive immune response in Arabidopsis *aba2-1* plants compared to WT and verified whether this could explain the lack of *Pst*-induced water-soaking lesions in *aba2-1* mutants. We started by investigating cROS production in response to a flg22 treatment in Arabidopsis *aba2-1* vs WT plants, in the presence or absence of DCMU. We found that Arabidopsis *aba2-1* mutant plants displayed constitutive cROS production, as well as apoplastic ROS (aROS) production, under control conditions, while both the WT and *aba2-1* mutant plants displayed similar cROS bursts in the presence of flg22 (Fig. 7, A and B). Interestingly, we observed that the constitutive cROS production in *aba2-1* plants was blocked by DCMU (Fig. 7, A and B).

**Figure 7.**
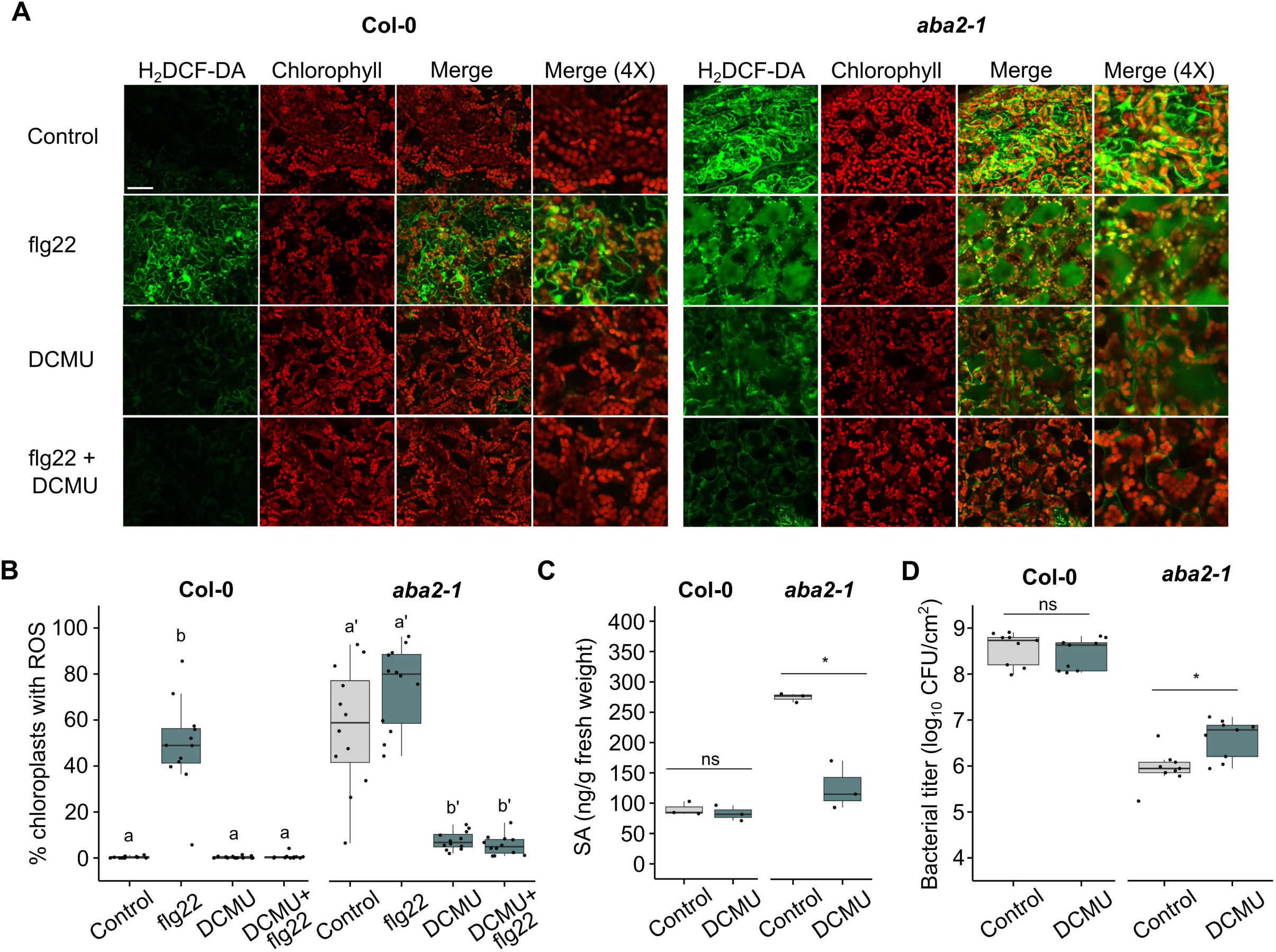
Constitutive cROS production and SA accumulation in the Arabidopsis aba2-1 mutant are not responsible for preventing the function of water-soaking effectors. (A) Leaves of four-wee-old WT and *aba2-1* Arabidopsis plants were syringe-infiltrated with either 10 mM MgCl_2_ (control), flg22 (1 μM), DCMU (10 μM), or a combination of flg22 and DCMU, as indicated. Five hours later, leaves were infiltrated with H_2_DCF-DA and leaf punches immediately harvested and mounted on microscopy slides, followed by visualization by confocal microscopy. Panels display channels detecting ROS production (H_2_DCF-DA), the presence of chloroplasts (chlorophyll), as well as merged images of the two (merge). Bars represent 50 μM. (B) Quantification of cROS detected in (A), presenting the number of cROS-positive chloroplasts as a percentage of total chloroplasts in images (% chloroplasts with ROS). Image analysis was performed by using the cell counter function of ImageJ after background removal. (C) Arabidopsis plants were treated as in (A), followed by quantification of salicylic acid (SA) by UPLC-MS at 24 hpt. (D) WT and *aba2-1* Arabidopsis plants were syringe-infiltrated with *Pst* WT (1 × 10^5^ CFU/ml), in the absence (control) or presence of DCMU (10 μM). Bacterial titers were quantified at 3 dpi. Statistical analyses were performed as followed. For 7D, a Wilcoxon-Mann-Whitney test was performed between the two treatments on Col-0, while a paired t-test was performed for *aba2-1* mutant plants. Different letters, or a * symbol, indicate statistical significance (p < 0.05), while ns = not significant.

We next questioned whether an abrogation of cROS bursts in *aba2-1* mutant plants would result in a reduction in SA accumulation. Arabidopsis WT and *aba2-1* plants were either sprayed with a mock solution, or with DCMU, 24 hours before tissues were harvested. Tissues were harvested at this timepoint to allow residual SA pools resulting from constitutive cROS production to be metabolized. At this timepoint, SA levels in DCMU-treated *aba2-1* plants resembled SA levels found in WT Arabidopsis control plants (Fig. 7C). We then asked whether cROS inhibition would result in differences in terms of bacterial load found in *aba2-1* mutants during an infection with *Pst*. We found that DCMU treatment resulted in a small, but statistically significant, increase in bacterial growth between Arabidopsis *aba2-1* mutant plants compared to controls, but not in WT plants subjected to the same treatment (Fig. 7D). This suggests that the constitutively produced cROS, and increased SA levels found in *aba2-1* plants only marginally contribute to disease resistance against *Pst*.

Why does inhibiting production of cROS and SA not lead to a greater increase in microbial growth? Since increased SA levels are able to prevent *Pst*-induced water-soaking^12^, we investigated whether *Pst* could disrupt cROS bursts in Arabidopsis *aba2-1* mutants as it does in WT plants to eventually suppress SA induction. Consistent with the other findings in this study, we found that *Pst* was unable to suppress cROS production in *aba2-1* mutants compared to WT plants (fig. S6A and S6B), suggesting that pathogen manipulation of ABA pathways is critical for preventing cROS bursts. Likewise, *Pst* inoculation resulted in enhanced SA accumulation in Arabidopsis *aba2-1* compared to WT plants (fig. S7A). However, DCMU treatment resulted in a total abrogation of SA accumulation above basal levels in both *aba2-1* and WT plants inoculated with *Pst* (fig. S7A). This provided us with a means to investigate whether cROS production, and therefore SA accumulation, in *aba2-1* mutants was responsible for the lack of *Pst*-induced water-soaking lesions. We hypothesized that although SA inhibits the ability of ABA to close stomata, SA levels are not sufficient to prevent *Pst*-induced water-soaking in *aba2-1* plants. Indeed, we found that *Pst* could not induce water-soaked lesions in *aba2*-*1* mutant plants that were co-treated with DCMU, which compromises SA accumulation (fig. S7B). This further supports the notion that *Pst* requires functional ABA pathways to induce a pathogenic niche in the apoplast^16,25^, and that increased SA accumulation alone is not sufficient to prevent water-soaking appearance in the *aba2-1* mutant. This is consistent with our previous suggestion that the major effect of a lack of ABA on a bacterial infection is that it precludes the ability to induce an aqueous apoplastic niche. Thus, our results are consistent with a role for cROS in inducing SA production, which under normal circumstances is required to counteract that stomatal closing effects of ABA. Thus, it is this effect that is ultimately a major contributor to defense against bacteria. However, in the absence of ABA, stomatal closure is not induced, even if SA production is compromised.

## Discussion

Chloroplasts are dynamic and critical organelles involved in a myriad of metabolic and physiological processes. Our study has revealed that ROS bursts generated from PSII following the perception of microbe/danger-associated peptides are indispensable for proper activation of the plant immune system. We demonstrate that inhibiting cROS production disrupts the production of SA without altering SA-independent functions of plant immunity, further supporting the hypothesis that cROS may be an initiator of SA biosynthesis in plant defense. We further show that cROS production, and photosynthesis more generally, are targeted by the conserved effectors HopM1 and AvrE1 in an ABA-dependent manner to reduce SA pools and increase *Pst* virulence. Understanding how pathogens disrupt host immunity, such as SA induction, could have critical implications in developing pathogen-resistant crops.

### Are cROS bursts at the origin of SA induction following immune activation?

Our results show a strong association between the induction of plant defenses, cROS production and SA production. In addition to plant immune activation, multiple stresses, such as photoperiod stress, induce the production of cROS^11^. Interestingly, photoperiod stress has been shown to induce SA accumulation, which blocks the ability of a bacterial pathogen to induce an aqueous apoplastic niche^12,14^. Similarly, salt and cold stress both lead to the induction of cROS, as well as the induction of SA-responsive genes and/or SA accumulation^18,20^. Long term high humidity treatment (over two days) has also been shown to induce cROS, as well as ethylene production and SA extracellular transport^26,27^. Whether cROS production is involved in SA biosynthesis in all of the above situations is unknown. However, a strong relation appears to exist between cROS and SA production beyond what is observed in plant defense responses. Another key factor, Ca^2+^, has also been shown to contribute to SA induction in the chloroplast^28^. Exploring whether, and how, chloroplastic Ca^2+^ is linked to SA induction by affecting cROS production will be of interest for future research.

### Why do pathogens target cROS production in plant immunity?

It has previously been reported that protecting plants by applying PAMPs prior to a disease challenge prevents the ability of a pathogen to suppress photosynthesis and cROS production^28^. It was proposed that *Pst* could disrupt photosynthesis, and potentially cROS bursts, by exploiting its host’s ABA pathway^15^. Manipulation of plant ABA pathways is a highly conserved virulence strategy used by plant pathogens to create an ideal extracellular niche containing water and nutrients^25,29^. ABA also antagonizes plant immune signaling, including SA-mediated immune responses. Given that the manipulation of the ABA pathway by *Pst* is mediated, in large part, by the effectors HopM1 and AvrE1^16,17^, it is logical that these effectors are also involved in the repression of SA accumulation via SA-ABA antagonism. We have previously shown that high levels of SA induced by photoperiod stress, or ectopic application of an SA analog, prevents pathogen-induced water-soaked lesions^12^. These observations raise the question of whether the impact of ABA alone on stomata is sufficient for the establishment of pathogenic niches, or whether the negative impact that ABA has on SA signaling also contributes to water-soaking. We observed that a DCMU treatment is sufficient to abrogate SA induction in Arabidopsis *aba2-1* mutant plants (Fig. 7C). However, this abrogation does not result in the appearance of water-soaked lesions (fig. S7B). Thus, our data suggest that the lack of water-soaking lesions observed in *aba2-1* plants^16,17^ is likely not caused directly by the constitutive SA accumulation found in this mutant line, but rather from the lack of ABA required to induce stomatal closure. However, we cannot rule out the possibility that the lack of water-soaking in *aba2-1* plants could be due to a constitutive SA accumulation throughout the plant’s life prior to the infection. Indeed, SA levels in *aba2-1* control plants are about twice as high as those found in WT control plants (Fig. 7C). Considering that plants displaying autoimmune phenotypes, such as the Arabidopsis *snc1* mutant, normally accumulate up to 10 times more SA than WT plants^30^, it is unlikely that the slightly enhanced SA accumulation found in Arabidopsis *aba2-1* mutant plants are sufficient to prevent water soaking. In fact, the levels of SA found in Arabidopsis *aba2-1* plants are similar to those observed in other Arabidopsis WT accessions in which *Pst* is still pathogenic^31^.

### How does ABA prevent SA biosynthesis in chloroplasts during plant immunity?

Previous studies have established that ABA inhibits photosynthesis, as well as the accumulation of SA^2,12,15,29^. It has been proposed that ABA globally impacts chloroplast functions and that the lack of SA induction in plant immunity could simply be the result of reduced chloroplastic activity. Supporting this hypothesis, ABA has been reported to negatively impact the total chloroplast transcription rate as well as reducing the carboxylase activity of Ribulose-1,6-bisphosphatase^32–34^. Our observation that the ABA-inducing effectors AvrE1 and HopM1 of *Pst* cause a global reduction in the transcription of genes related to chloroplast function are consistent with such a global effect of ABA (Figure 5E, 5F). It has also been proposed that the decrease in substomatal CO_2_ concentration caused by ABA-induced stomatal closure leads to a global reduction in photosynthesis and thus, potentially less cROS induction during immune elicitation. Which of these hypotheses is the lead cause behind ABA-induced photosynthesis reduction has long been debated and it is possible that both mechanisms are involved in ABA-mediated suppression of SA accumulation.

It is interesting to note the complete disruption of SA accumulation during a co-treatment of the PAMP flg22 with the PSII inhibitor DCMU (Fig. 4C), but not during an infection with a virulent *Pst* strain (Fig. 6D), which suppresses cROS bursts and reduces photosynthesis. It is likely that the specificity with which DCMU targets PSII activity may function more potently than pathogen-induced ABA itself. Also, syringe-infiltration of DCMU alongside an immunogenic peptide assures that, most likely, all cells in contact with the solution will rapidly experience reduced photosynthesis and cROS production. In contrast, it is possible that not all cells undergo cROS inhibition upon syringe-infiltration of bacteria, since effectors are likely not delivered homogenously to all cells^35,36^. This would allow some cells to still respond to the pathogen and produce SA, as has been observed in plant cells bordering important apoplastic bacterial populations^37^. On the other hand, since *Pst* effectors affect multiple phytohormones during an infection^2^, it is possible that in addition to cell-surface pathogen perception, SA accumulation may also be induced by the impact of effectors on host metabolic processes. Although speculative, this would suggest that plants may have developed a strategy to monitor their metabolism following an infection, thus permitting SA induction in the absence of cROS. Another explanation for the presence of SA during *Pst* infection may be due to the detection limit of the ROS probe H_2_DCF-DA during an infection, in which case cROS may simply be incompletely repressed by HopM1 and AvrE1. Finally, it is also possible that cROS generated from PSII mechanisms are not the sole contributors to SA induction during an infection process and that these other pathways are not as efficiently targeted by *Pst* effectors. Together, these results suggest that, although other mechanisms may exist, cROS are a major signal for the induction of SA production.

Further implications of cROS burst in plant immunity may be to remodel chloroplast morphology^10^, such as by inducing stromule formation^6,7^, which has been hypothesized to act in chloroplast-to-nucleus retrograde signaling. Interestingly, stromules have been shown to contribute to plant defense and are induced by immunogenic compounds, including flg22, as well as by SA and H_2_O_2_^6^. A more complete model appears to emerge from this and other studies, in which pathogen perception leads to cROS production, which alters chloroplast morphology and function, including the induction of SA biosynthesis to fend off potential invaders. The establishment of cROS production as an anchor point in the initiation of SA production raises multiple additional questions such as how, for example, PAMP perception at the cell surface leads to changes in the chloroplast. Given that the latter take place somewhat later than early signaling events (Figure 1, 3) this implies a series of as yet unknown signaling events that affect chloroplast metabolism. Our results indicate that defense-induced cROS are a byproduct of photosynthesis. However, the mechanism leading to an increase in ROS production remains to be defined. Likewise, the immediate effects of cROS in modulating enzyme function and/or gene regulation remain to be explored.

In conclusion, our data provide a link between cROS production and the induction of SA in plant immunity, with cROS emerging as a strong candidate for being the initiator of a SA-mediated plant immune responses. Furthermore, this study sheds light on a commonly observed physiological impact of microbial infection, that is, a reduction in host photosynthetic functions, as well as a hypothesis for why this phenomenon is so widely conserved.

## Authors contributions

C.R.-L. designed the study. C.R.L. and P.M. supervised the study. C.R.-L., M.St-A., P.D.-F., R.P., A.P., S.G., and F.G.-L. performed the experiments. C.R.-L. and M.St-A. performed bioinformatic analysis. C.R.-L. and R.P. analyzed the data and generated the figures. C.R.-L. wrote the manuscript. C.R.-L., I.L.-L. and P.M. edited the manuscript. P.M. and I.L.-L. secured funding.

## Acknowledgements

The authors would like to thank Jacqueline Monaghan, Alexander M. Jones, Bijun Tang, Sheng Yang He, as well as Jong-Hum Kim, for the great discussions regarding the findings in this study. The authors would also like to thank Christian Danve Marco Castroverde for critical reading of the manuscript and his inputs for the study more broadly. This study was supported by two Natural Sciences and Engineering Research Council of Canada (NSERC) Discovery Grants respectively to P.M. and I.L.-L., as well as by a Fonds de Recherche du Québec—Nature et Technologies (FRQ-NT) Team Grant and NOVA-NSERC Alliance Grant to P.M. (PI) and I.L.-L. (co-PI). I.L.-L. is also supported by a Canada Research Chair T2. C.R.-L. was supported by a VoiceAge Excellence Scholarship from the Faculty of Science of Université de Sherbrooke and by a Quebec Doctoral Graduate Scholarship (FRQ-NT). A.P. was supported by a Canada’s Master’s Graduate Scholarship (NSERC). R.P. was supported by a Quebec Master’s Degree Graduate Scholarship (FRQ-NT).

## Material and methods

### Plant material

Arabidopsis plants were grown in PromixTM soil (PremierTech) in growth chambers with a 12 h light/dark photoperiod, with relative humidity of ∼60% at 21 °C. Light intensity was measured at 180 µmoles/m^2^/s when lights were on. Four-to five-week-old Arabidopsis plants were used for all experiments described herein, unless stated otherwise.

### Bacterial disease assays

*Pseudomonas syringae pv. tomato* DC3000 WT and mutant strains were cultured overnight at 28 °C in Luria-Bertani (LB) media containing 50 mg/l of rifampicin. On the day of the infection, fresh LB media was inoculated with 0.5 ml of the overnight culture and bacteria were collected when OD_600_ reached between 0.8–1. Bacteria were centrifuged for 10 min at 4000 × g and the pellet resuspended in MgCl_2_ 10 mM. Bacterial density was adjusted to 0.2 (1 × 10^8^ CFU/ml) prior to further dilutions. Bacterial infections were carried out between 14:00–15:00 (zeitgeber time of 6:00–7:00).

All inoculations of Arabidopsis leaves were performed by syringe-infiltration. Infiltrated plants were all kept under ambient humidity levels for 1–2 h to allow water to evaporate, then domed with a plastic unit to maintain high humidity (>95% RH).

Bacterial growth *in planta* was monitored by harvesting infected Arabidopsis leaves, surface sterilizing in 80% ethanol and rinsing in sterile water twice. Leaf disks were taken from three leaves from the same plant (one per leaf; total of three leaf disks) using a cork borer (6 mm in diameter) and ground in sterile 10 mM MgCl_2_. Three biological replicates were performed for each experiment. Colony-forming units (CFU) were determined by making serial dilutions (10^0^−10^−6^) and plating on LB plates containing 50 mg/l of rifampicin. Each dilution was plated in three technical replicates. Experiments were repeated at least three times and values from replicates are tabulated in the Source Data File.

### Chemical treatments

To assess plant immune responses, flg22, elf18 or AtPep1 were synthesized (BioBasic) and syringe-infiltrated (1 μM) into Arabidopsis leaves. To disrupt host photosynthetic functions, plants were treated with 3-(3,4-dichlorophenyl)-1,1-dimethylurea (DCMU; Millipore-Sigma). To assess plant responses salicylic acid (SA; Millipore-Sigma), leaves were sprayed with a 0.5 μM SA solution.

### ROS quantification

For apoplastic ROS quantification, leaf disks from four-week-old Arabidopsis plants were collected using a 4 mm diameter biopsy punch and placed into white 96-well plates (Corning) containing 100 μl of distilled water for 16 h (overnight). Prior to ROS quantification, water was removed and replaced with ROS assay solution (100 μM Luminol [Millipore-Sigma], 20 μg/ml horseradish peroxidase [Millipore-Sigma]), with or without immune elicitors. Light emission was measured using a TECAN Spark® plate reader. Experiments were repeated three times and values from replicates are tabulated in the Source Data File.

For chloroplast ROS imaging, confocal microscopy was performed as previously described^15^, with modifications. Leaves were syringe-infiltrated with control or bacterial solutions, biotic stressors, or compounds, and blotted with tissue paper to remove excess liquid. For PAMP/DAMP and DCMU treatment, plants were returned to growth chambers for the time indicated in individual figure legends. In our hands, cROS could be observed with high consistency starting at 4 hours post-treatment (hpt) with PAMP/DAMP. After 5-6 hpt, leaves were syringe-infiltrated with 10 μM of 2′7′-dichlorodihydrofluorescein diacetate (H_2_DCF-DA; Millipore-Sigma) and blotted dry. A section of the leaf (avoiding the site damaged by the syringe) was excised with a razor blade and mounted in perfluorodecalin (Millipore-Sigma). Microscopy slides were imaged within 30 minutes following mounting on the microscopy slide. Images were acquired using a FV-3000 Olympus confocal microscope. Images were analyzed and chloroplasts positive for ROS production were quantified using ImageJ software.

### Callose deposition imaging and quantification

Leaves of four-week-old Arabidopsis leaves were infiltrated with *Pst hrcC^-^* (1 x 10^8^ CFU/ml), or other indicated compounds, 24 hours prior to tissue harvesting for callose staining. Leaf disks were taken from four leaves from the same plant (one per leaf; total of four-leaf disks) using a cork borer (6 mm in diameter) and placed in 24-well plates containing 95% ethanol. Plates were placed on a rotating shaker and ethanol renewed three times over the course of four hours to allow leaf decolouration. Once leaf disks were decoloured (white), they were rinsed in 67 mM K_2_HPO_4_ for 1 hour. K_2_HPO_4_ was removed and replaced by callose staining solution (K_2_HPO_4_ 67mM containing 0.1% aniline blue) for another hour. The staining solution was then removed, and leaf-disks quickly rinsed in 67 mM K_2_HPO_4_. The rinsing solution was then removed and replaced with fresh 67 mM K_2_HPO_4_ solution and placed back on the rotating shaker. The plate containing leaf disks was then placed in a 4°C refrigerator until imaging.

Callose deposition was imaged using a FV-3000 Olympus confocal microscope. Callose quantification was measured using a reversed-color analysis function of ImageJ software and quantifying the amount of callose deposition in terms of pixels against the total amount of pixels found in each image, which resulted in a percentage of callose area per image.

### RNA extraction and real-time quantitative PCR

RNA was extracted from frozen and ground leaf tissue using QIAZOL (QIAGEN) reagents, followed by on-column DNase treatment (QIAGEN), according to the manufacturer’s protocol. RNA purity was assessed with a spectrophotometer and quality by gel electrophoresis. cDNA was generated by using MMuLV-RT (Service de purification des proteins, Université de Sherbrooke).

Quantitative real-time PCR was performed with a Bio-Rad CFX96 machine. Each reaction contained 1X SYBR mix (Service de purification des proteins, Université de Sherbrooke), specific primers and a 1:20 dilution of 500 ng of cDNA stock. Amplification cycle protocols were as follow: 2 min at 95 °C; 40 cycles of 6 s at 95 °C and 30 s at 60 °C. Melting curves were verified at the end of 40 cycles for confirmation of primer specificities. All reactions were repeated in three technical and biological replicates. Average Cq values were normalized by ΔΔCT formula against *ACTIN2* expression. Oligonucleotide primers used in this study can be found in Supplementary Table 1.

### Salicylic acid quantification by UPLC-MS

Leaves of four-week-old Arabidopsis plants were harvested and weighed for fresh weight calculation and immediately flash-freeze in liquid nitrogen. Tissues were ground with a plastic pestle and phytohormones extracted overnight using 0.5-1 ml of ice-cold extraction buffer (methanol: water (80:20 v/v), 0.1% formic acid, 0.1 g/L butylated hydroxytoluene and 100 nM ABA-d6 as an internal standard), as described previously^12,16^. Extracted phytohormones were filtered by using several rounds of centrifugation and supernatant collection.

Filtered extracts were quantified using an Acquity Ultra Performance Liquid Chromatography system (Waters Corporation, Milford, MA) as described previously^16^. SA was quantified based on a standard curve to calculate sample concentration (nM), which was converted to ng using the molecular weight of each specific compound and the extraction volume used. All data were normalized to initial fresh weight in grams. Experiments were repeated three times and values from replicates are tabulated in the Source Data File.

### RNA-sequencing analysis

For RNA-sequencing analysis, data were obtained from a previous study^16^, which have been deposited at GEO (Gene Expression Omnibus) under the accession number GSE186836. Briefly, the data previously obtained from this study were re-analyzed for the GO terms with significant down-regulation compared to control plants. GO term analysis was performed with the R package for PANTHER/REVIGO^38^. Heatmaps represent log2-fold relative expression compared to control plants (infiltrated with 10 mM MgCl_2_) and have been generated using the R package pheatmap^39^.

### Deep-phenotyping and photosynthetic parameters analysis

For plant phenotyping and photosynthetic parameters analysis, plants were transferred from growth chambers to a PlantScreen™ phenotyping system (Photon Systems Instruments, Drásov, Czechia) at the Eastern Canadian Plant Phenotyping Platform (Sherbrooke). First, RGB images were acquired using a plant screen to isolate plant-associated pixels from background pixels. For *F_v_/F_m_* analysis, plants were dark-adapted for 15 minutes prior to starting the *F_v_/F_m_* protocol, according to the manufacturer’s instructions. NPQ analysis was performed concomitantly with the *F_v_/F_m_* protocol, according to the manufacturer’s instructions.

### Statistical analysis

All statistical analyses were performed using the bioinformatic software R. Statistical significance was set at p<0.05. Statistical tests used are stated in figure legends. All statistical test assumptions, such as normality and homoskedasticity, were tested. When not respected, non-parametric equivalent tests were performed. In multiple comparison tests, Bonferroni corrections were used. Data points represented in chloroplast ROS bursts analysis represent a percentage of chloroplast in one microscopy image, which generally consists of between 300-900 chloroplasts. Chloroplast figures originated from at least three leaves from three different plants.

## Supplementary figures

**Figure S1 | Impact of photosynthesis inhibition on ETI**

(A) One half of leaves of four-week-old Arabidopsis plants were infiltrated with either 10 mM MgCl_2_ (control), DCMU (10 μM), *Pst* DC3000 or DC3000 expressing AvrRpm1 or AvrRps4 (1 × 10^8^ CFU/ml), in the presence or absence of DCMU, as indicated. Infiltrated leaves were photographed at 12 hours post infiltration. (B) Arabidopsis plants were syringe-infiltrated with *Pst* AvrRpm1 or *Pst* AvrRps4 (1 × 10^5^ CFU/ml), in the absence or presence of DCMU (10 μM), as indicated. Bacterial titers were quantified at 3 dpi. A Student’s t-test was performed in S1B. A * symbol indicates statistical significance (p < 0.05).

**Figure S2 | Salicylic acid does not trigger cROS production**

(A) Leaves of four-week-old Arabidopsis WT or *ics1* mutant plants were syringe-infiltrated with either 10 mM MgCl_2_ (control) or flg22 (1 μM). Five hours later, leaves were infiltrated with H_2_DCF-DA and leaf punches immediately harvested and mounted on microscopy slides, followed by visualization by confocal microscopy. Panels display channels detecting ROS production (H_2_DCF-DA), the presence of chloroplasts (chlorophyll), as well as merged images of the two (merge). Bars represent 50 μM. (B) Quantification of cROS detected in (A), presenting the number of cROS-positive chloroplasts as a percentage of total chloroplasts in images (% chloroplasts with ROS). (C) Leaves of four-week-old Arabidopsis WT or *ics1* mutant plant leaves were syringe-infiltrated with either 10 mM MgCl_2_ (control), flg22 (1 μM) or SA (0.5 μM) and treated as in (A). Panels are displayed as in (A). (D) Quantification of cROS detected in (C), presenting the number of cROS-positive chloroplasts as a percentage of total chloroplasts in images (% chloroplasts with ROS). Image analysis was performed by using the cell counter function of ImageJ after background removal. All experiments were repeated at least three times with similar outcomes. Statistical analyses were performed as follows: in B, a Wilcoxon-Mann-Whitney test was performed to compare the different treatment between Col-0 and *ics1*, respectively. In D, a Kruskal-Wallis test was performed to compare the different treatments. Different letters indicate statistical significance, while ns = not significant.

**Figure S3 | Impact of HopM1 and AvrE1 delivery of the expression of chloroplastic genes** Heatmap representing expression differences of genes encoded in the Arabidopsis chloroplast genome 36 hours after inoculation with the indicated *Pst* strains (1 × 10^6^ CFU/ml).

**Figure S4 | Impact of photosynthesis inhibition on effector-induced water-soaking lesions and disease development**

(A) Leaves of four-week-old Arabidopsis plants were syringe-infiltrated with either *Pst* DC3000 or *Pst h^-^/a^-^*(1 × 10^8^ CFU/ml) in the absence (control) or presence of 10 μM DCMU, as indicated. Leaves were photographed 24 hours post syringe infiltration. (B) Leaves of four-week-old Arabidopsis plants were syringe-infiltrated with either *Pst* DC3000 or *Pst h^-^/a^-^* (1 × 10^5^ CFU/ml), in the absence (control) or presence of 10 μM DCMU, as indicated. Bacterial titers were quantified at 3 dpi. A Student’s t-test was performed in S5B. A * symbol indicates statistical significance (p < 0.05), ns = not significant.

**Figure S5 | Photosynthesis inhibition allows for the development of water-soaked lesions under constant light**

(A) Leaves of four-week-old Arabidopsis plants were syringe-infiltrated with either 10 mM MgCl_2_ (control) or *Pst* DC3000 (1 × 10^8^ CFU/ml) in the absence (-) or presence (+) of 10 μM DCMU, as indicated. Plants were subsequently placed under a constant light (CL) regime of 24 hours light or kept under a regular light (RL) regime of 12 hours light/dark. Leaves were photographed 24 hours post syringe infiltration. (B) Leaves of Arabidopsis plants were syringe infiltrated with *Pst* DC3000 (1 × 10^5^ CFU/ml) in the absence (control) or presence of 10 μM DCMU, followed by growth under the light regimes described in (A). Bacterial titers were quantified at 3 dpi. A Student’s t-test was performed in S6B. A * symbol indicates statistical significance (p < 0.05)

**Figure S6 | Pst DC3000 requires plant ABA biosynthesis to suppress cROS production**

(A) Leaves of four-week-old wild-type (Col-0) or *aba2-1* mutant Arabidopsis plants were syringe-infiltrated with either 10 mM MgCl_2_ (control) or 1 × 10^8^ CFU/ml *Pst* (DC3000). Six hours later, leaves were infiltrated with H_2_DCF-DA and leaf punches immediately harvested and mounted on microscopy slides, followed by visualization by confocal microscopy. Panels display channels detecting ROS production (H_2_DCF-DA), the presence of chloroplasts (chlorophyll), as well as merged images of the two (merge). Bars represent 50 μM. (B) Quantification of cROS detected in (A), presenting the number of cROS-positive chloroplasts as a percentage of total chloroplasts in images (% chloroplasts with ROS). Image analysis was performed by using the cell counter function of ImageJ after background removal. A Student’s t-test was performed in S7B. ns = not significant.

**Figure S7 | Photosynthesis inhibition does not restore Pst-mediated water-soaking lesions in the Arabidopsis aba2-1 mutant**

(A) Leaves of four-week-old Col-0 and *aba2-1* Arabidopsis plants were infiltrated with 10 mM MgCl_2_ (control), 1 × 10^8^ CFU/ml *Pst* (DC3000), DCMU (10 μM) or a combination of *Pst* and DCMU. Leaves were harvested at 24 hpt and salicylic acid (SA) was quantified by UPLC-MS. A Kruskal-Wallis test was performed to compare the different treatments. Different letters indicate statistical significance. (B) Leaves of four-week-old Col-0 or *aba2-1* Arabidopsis plants were syringe-infiltrated with *Pst* DC3000 (1 × 10^8^ CFU/ml), in a solution containing either 10 mM MgCl_2_ (control) or 10 mM MgCl_2_ with an addition of 10 μM DCMU, as indicated. Leaves were photographed at 24 hpi.

