## Supplementary figures for "Chloroplastic ROS bursts initiate salicylic acid biosynthesis in plant immunity"

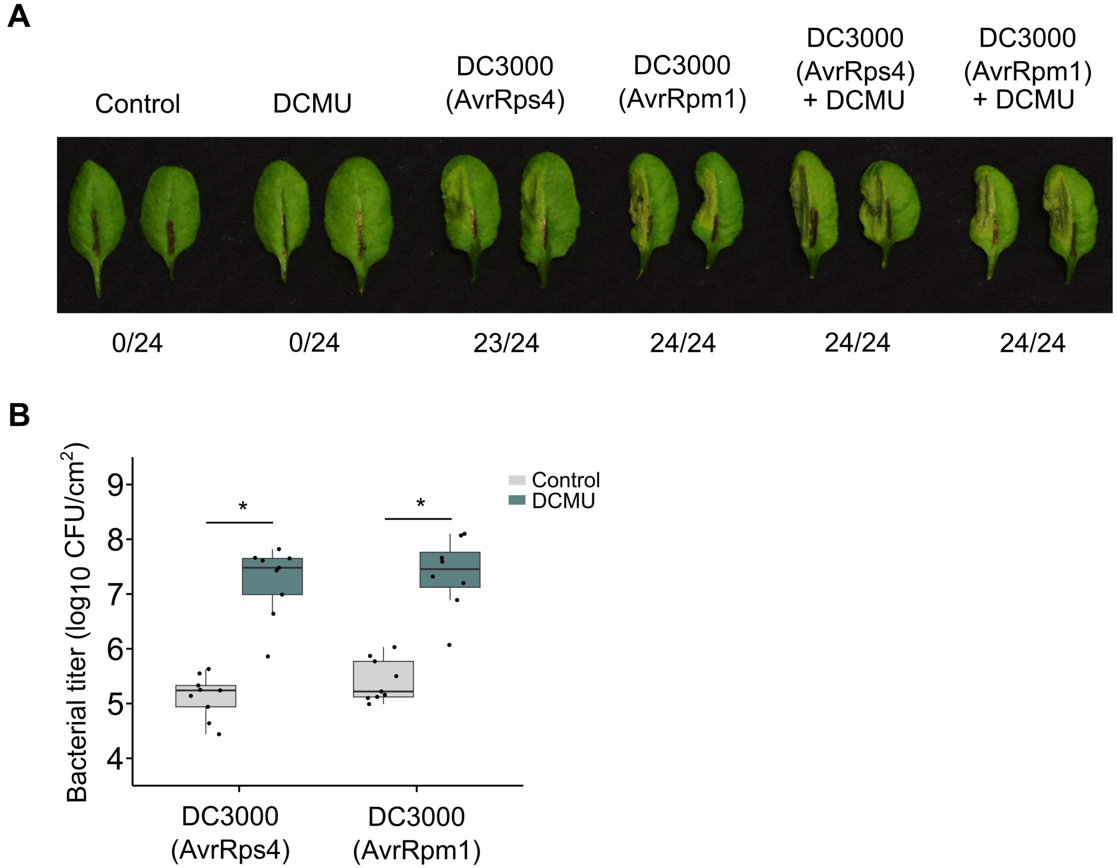


**Supplementary figure 1 | Impact of photosynthesis inhibition on effector-triggered immunity**

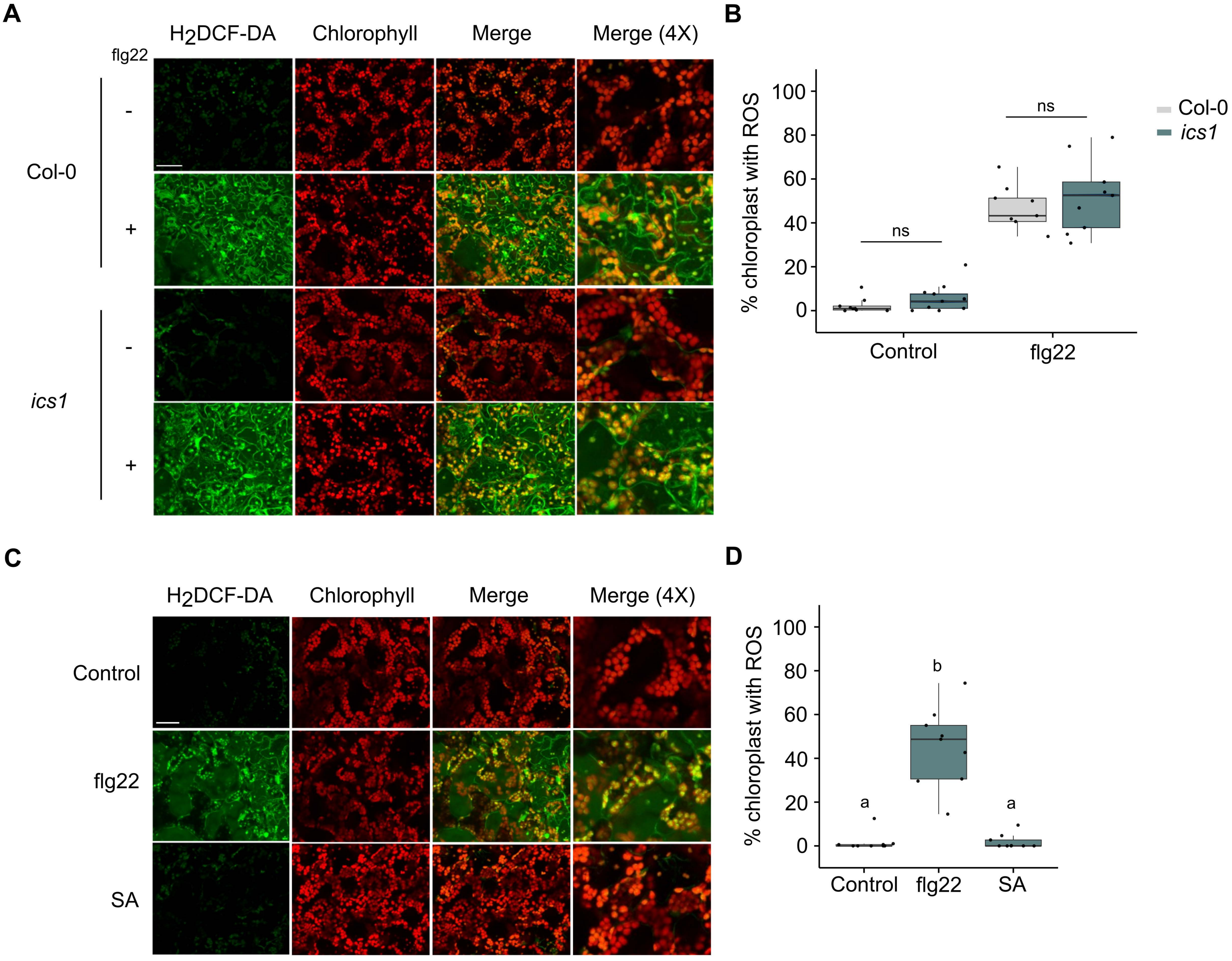


**Supplementary figure 2 | Salicylic acid does not trigger cROS production**

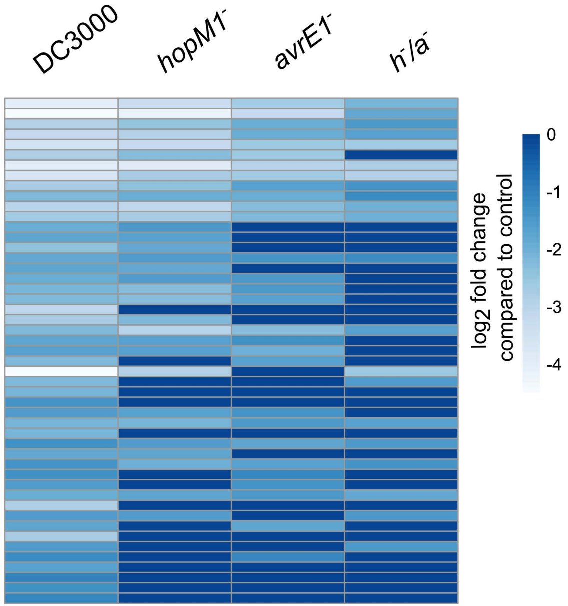


**Supplementary figure 3 | Impact of HopM1 and AvrE1 delivery of the expression of chloroplastic genes**

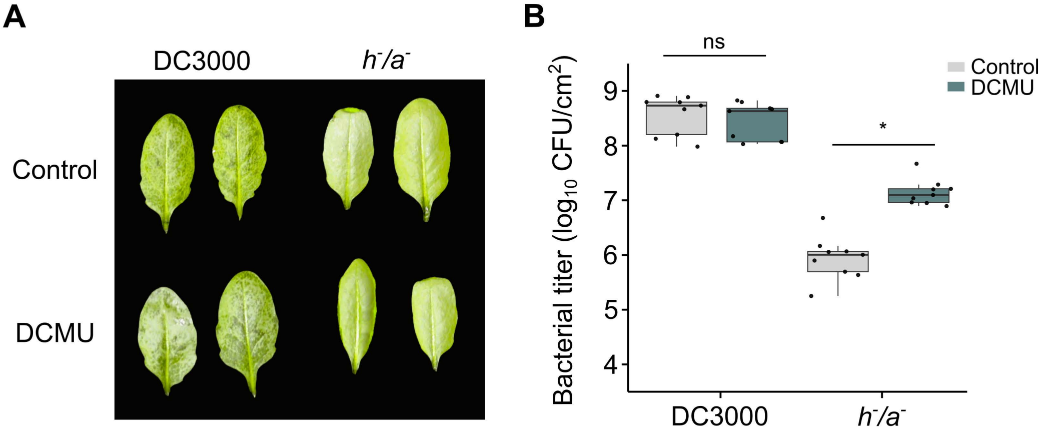


**Supplementary figure 4 | Impact of photosynthesis inhibition on effector-induced water-soaking lesions and disease development**

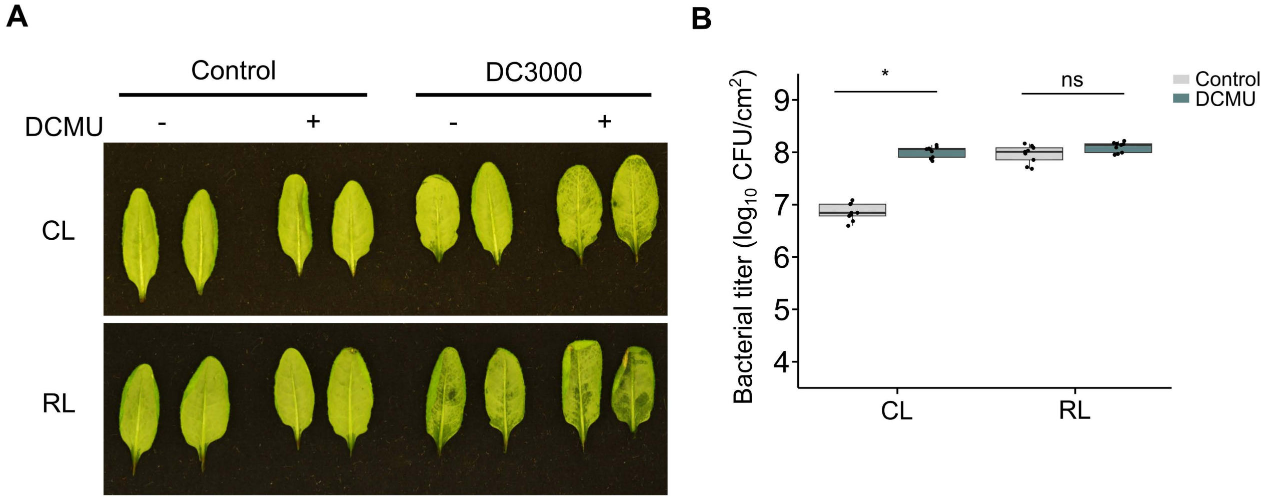


**Supplementary figure 5 | Photosynthesis inhibition allows for the development of water-soaked lesions under constant light**

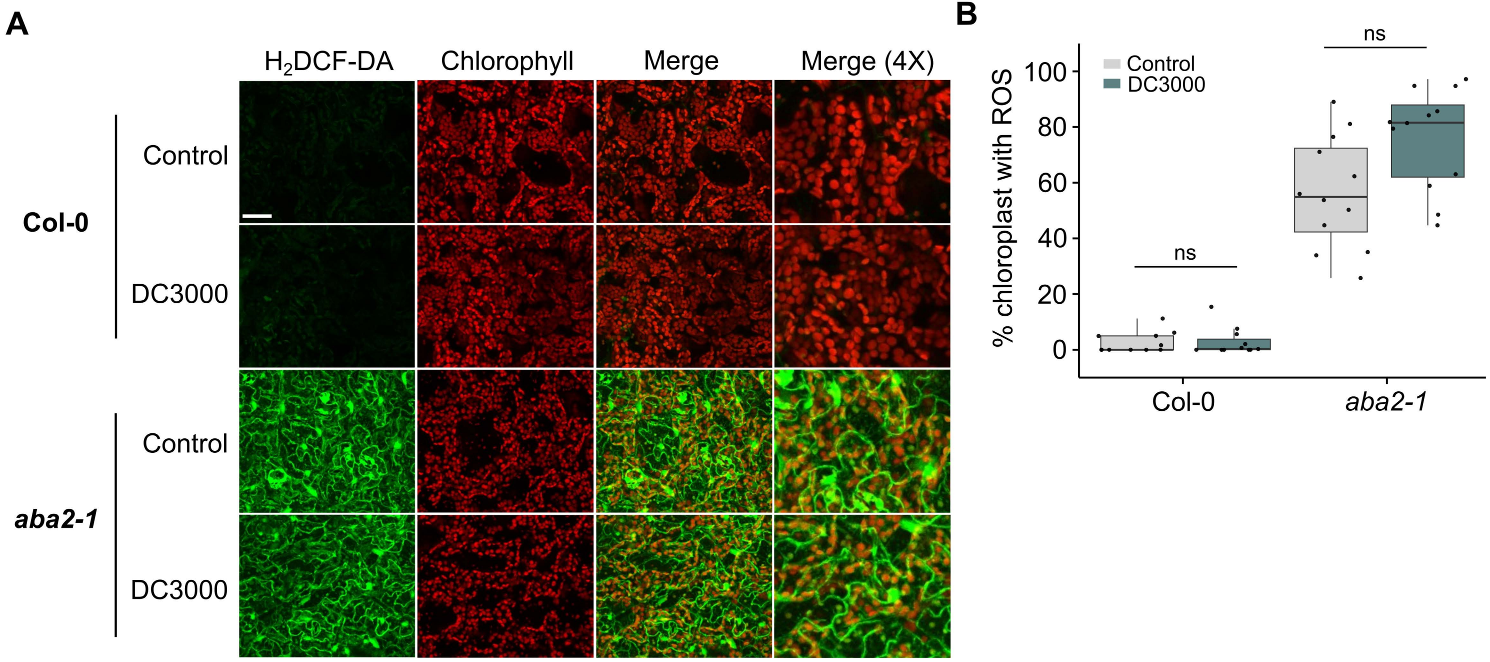


**Supplementary figure 6 | Pst requires plant ABA biosynthesis to suppress cROS production**

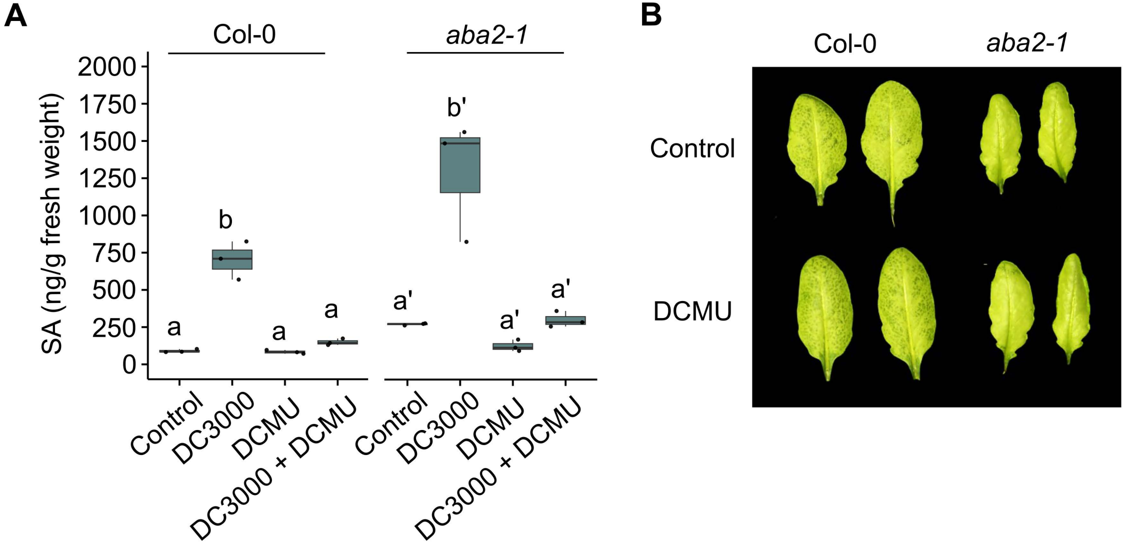


**Supplementary figure 7 | Photosynthesis inhibition does not restore *Pst*-mediated water-soaking lesions in the Arabidopsis *aba2-1* mutant**
